# Juvenile hormone interacts with multiple factors to modulate aggression and dominance in groups of orphan bumble bee (*Bombus terrestris*) workers

**DOI:** 10.1101/626382

**Authors:** Atul Pandey, Uzi Motro, Guy Bloch

**Affiliations:** Department of Ecology, Evolution and Behavior, The Hebrew University of Jerusalem, Israel

**Keywords:** Juvenile hormone, bumble bee, Bombus terrestris, dominance, aggression, social behavior, eusociality, reproduction, division of labor

## Abstract

Juvenile hormone (JH) is a key regulator of insect development and reproduction. Given that JH commonly affects adult insect fertility, it has been hypothesized to also regulate behaviors such as dominance and aggression that are associated with reproduction. We tested this hypothesis in the bumble bee *Bombus terrestris* for which JH has been shown to be the major gonadotropin. We used the allatoxin Precocene-I (P-I) to reduce hemolymph JH titers and replacement therapy with the natural JH to revert this effect. In small orphan groups of workers with similar body size but mixed treatment, P-I treated bees showed lower aggressiveness, oogenesis, and dominance rank compared with control and replacement therapy treated bees. In similar groups in which all bees were treated similarly, there was a clear dominance hierarchy, even in P-I and replacement therapy treatment groups in which the bees showed similar levels of ovarian activation. In a similar experiment in which bees differed in body size, larger bees were more likely to be dominant despite their similar JH treatment and ovarian state. In the last experiment, we show that JH manipulation does not affect dominance rank in groups that had already established a stable dominance hierarchy. These findings solve previous ambiguities concerning whether or not JH affects dominance in bumble bees. JH positively affects dominance, but bees with similar levels of JH can nevertheless establish dominance hierarchies. Thus, multiple factors including JH, body size, and previous experience affect dominance and aggression in social bumble bees.

## 1. Introduction

Juvenile hormone (JH) is a well-studied hormone named after its morphogenic functions in the regulation of insect metamorphosis. In adults of many, but not all, insects studied so far, JH functions as a gonadotropic hormone regulating fertility (Adams, 2009; De Loof et al., 2001; Riddiford, 2012). One of the best studied examples for a non-gonadotropin function in adult insects is the Western honeybee (*Apis mellifera*), in which JH does not affect adult female oogenesis but rather regulates age-related division of labor (Reviewed in Bloch et al., 2002; Hartfelder, 2000; Robinson and Vargo, 1997; Wegener et al., 2013). These findings for the honeybee contrast with evidence that JH retains its ancestral gonadotropic functions in species of bees and wasps that live in solitary lifestyle or in small and simple societies (Bell, 1973; Bloch et al., 2000a, 1996; Chen et al., 1979; Pan and Wyatt, 1971; Shorter and Tibbetts, 2009; Shpigler et al., 2014; Smith et al., 2013; Wasielewski et al., 2011). These observations led to hypotheses stating that the evolution of advanced sociality in bees was associated with modification in JH signaling pathways (Bloch et. al., 2002; Hartfelder K, 1998; Robinson and Vargo, 1997; West-Eberhard, 1996). There is also evidence consistent with the premise that JH lost its ancestral gonadotrophic functions along the evolution of advanced sociality in additional lineages such as ants and wasps (Giray et al., 2005; Lengyel et al., 2007; Norman and Hughes, 2016; O’Donnell and Jeanne, 1993; Penick et al., 2011; Shorter and Tibbetts, 2009). This association between modifications in JH functions and the evolution of sociality is commonly attributed to assumed fitness costs related to high JH titers, similar to the well-established costs of testosterone in vertebrates (Flatt and Kawecki, 2007; Rantala et al., 2003; Rodrigues and Flatt, 2016). Accordingly, the loss of JH gonadotropic functions in complex insect societies is assumed to allow highly social insect queens to be exceptionally fertile over long periods while escaping the hypothesized fitness cost of high JH levels (Pamminger et al., 2016; Rodrigues and Flatt, 2016).

Bumble bees provide an excellent model system for studying proximate and ultimate aspects of the relationships between JH signaling and sociality because they are social, but their societies are in many aspects simpler than that of the highly eusocial honeybees and ants. By contrast to honeybees, JH does not influence task performance (Shpigler et al., 2016), and there is good evidence consistent with the hypothesis that JH is the major gonadotropin in bumble bees. JH titers are positively correlated with ovarian activity (Bloch et al., 2000a, 1996), and manipulations increasing or decreasing JH levels, result in ovarian activation or inactivation, respectively (Amsalem et al., 2014b; Röseler, 1977; Shpigler et al., 2014, 2010, 2016; Van Doorn, 1989). JH was also shown to regulate additional processes that are associated with reproduction such as wax secretion and comb construction (Shpigler et al., 2014). As expected for a gonadotropic hormone, JH treatment upregulates the expression of the major yolk protein vitellogenin (Vg) in the fat body and its protein levels in the hemolymph (Shpigler et al., 2014).

In vertebrates such as mammals, birds, and fish, the gonadotropic steroid hormones commonly regulate behaviors related to reproduction such as courtship, mating, aggression, and dominance (Brain, 1977; Nelson, 2005; Norris and Lopez, 2011). The relationships between gonadotropic hormones and behavior have received significantly less attention in insects and other invertebrates. The current study aims at testing the hypothesis that JH regulates agonistic behaviors underlying dominance hierarchy establishment in the social bumble bee *Bombus terrestris*.

Dominance rank is typically achieved by means of overt aggression and agonistic interactions and is an important factor determining reproduction in many eusocial insects, including bumble bees (Andrade-Silva and Nascimento, 2015; Bloch et al., 1996; Duchateau and Velthuis, 1989; Geva et al., 2005; Monnin and Peeters, 1999; Roseler, 1991; Sasaki et al., 2016; Van Doorn, 1989; Van Doorn and Heringa, 1986). Earlier studies on JH, dominance, and aggression in bumble bees led to somewhat conflicting conclusions. In *B. terrestris*, the best-studied bumble bee, JH titers, and in-vitro biosynthesis rates are typically correlated with dominance rank; dominant queens and workers have active ovaries, large corpora allata (the JH producing glands) and high JH titers; dominant workers also typically show more overt aggression and threatening displays (Amsalem et al., 2014b; Amsalem and Hefetz, 2011; Bloch et al., 2010, 2000a; Larrere and Couillaud, 1993; Röseler, 1977; Van Doorn, 1989). However, in small groups of callow workers, a dominance hierarchy is established and aggression is typically highest during the first few days post pupal emergence, a period when hemolymph JH titers are still low. At later ages when the dominance hierarchy is already clear, some subordinate individuals, which show little aggression or dominance displays, may nevertheless have developed ovaries or high JH titers (Bloch et al., 2000a, 1996; Röseler, 1977; Van Doorn, 1989; Van Doorn and Heringa, 1986). A recent study in which circulating JH titers were reduced using the allatotoxin precocene-I (P-I), reported reduced aggressive behavior, but the interpretation of this finding is difficult because the effect could not be recovered by replacement therapy with JH-III (Amsalem et al., 2014b).

To thoroughly test the hypothesis that JH influences aggression and dominance, we manipulated circulating JH titers in callow orphan (“queenless”) workers using a combination of JH reduction with P-I, and replacement therapy with JH-III, the natural JH of bumble bees (Bloch et al., 2000, 1996). We tested the hypothesis that JH influence dominance and aggression in bumble bee in 4 complementary experiments: In Experiment 1, we studied queenless groups composed of same body size, same age workers, each subjected to a different JH treatment. This experiment directly tested whether a decrease or an increase in JH levels affect dominance rank. In Experiment 2, we studied similar groups composed of workers of different body size, but of similar age, and subjected to the same treatment. This experiment tested whether body size influences dominance hierarchy in bees with similar JH levels. In Experiment 3, we studied groups of workers with a similar body size, age and subjected to the same JH treatment. This experiment tested whether bees with similar JH levels and size nevertheless establish dominance hierarchies. In the last experiment (Experiment 4), we studied queenless groups which had already established a dominance hierarchy and tested whether treatment with P-I or JH can reduce or increase, the dominance rank of the most dominant or most subordinate individual, respectively. This experiment tested the interaction between JH levels and previous experience (i.e., dominance rank). Taken together, our results suggest that JH is one of several factors affecting aggressiveness and dominance behavior in bumble bees and can explain the inconsistent results in previous studies on JH and dominance in bumble bees.

## 2. Materials and Methods

### 2.1. Bees

*Bombus terrestris* colonies with a queen, 5-10 workers and brood at various stages of development (typically 2–4 days post first worker emergence) were obtained from Polyam Pollination Services, Yad-Mordechai, Israel. Each colony was housed in a wooden nesting box (21 × 21 × 12 cm) and placed in an environmental chamber (29 ± 1 °C; 55% ± 10% RH) in constant darkness at the Bee Research Facility at the Edmond J. Safra campus of the Hebrew University of Jerusalem, Givat Ram, Jerusalem. The top and front walls of the nest boxes were made of transparent Plexiglass panels, and the other parts were made of wood. Commercial sugar syrup (Polyam Pollination Services) and fresh pollen (collected by honey bees; mixed with sugar syrup) were provided *ad libitum*. We painted each individual bee within the treatment with xylene free silver color paint (Pilot-PL01735) on the thorax creating a similar pattern across treatment groups. All treatments and behavioral observations were performed under dim red light. Given that bumble bees are very sensitive to substrate-borne vibrations, we paid special attention to minimize touching the cages or substrate during treatments and observations.

### 2.2. Manipulating JH titers

Our protocols are based on those developed and validated by Shpigler et al., (2016) with the following modifications: (A) We kept the concentration of P-I constant at 50µg/µl and varied the volume in which it was dissolved in order to adjust the total amount applied to a bee to her body-size (see below). (B) In the second treatment, we applied the JH-III or vehicle to the abdomen rather than to the thorax. This was done in order to minimize possible mixing of the P-I and JH treatments, and reduce the number of times the bee is anesthetized by chilling. (C) We used a higher and somewhat different range of doses (200-260μg/bee compared to 160-210μg/bee). In brief, we collected callow workers (<24 h after emergence from the pupa) and cold anesthetized them in 50 ml tubes immersed in ice (∼2 °C) until being immobile for 5– 10 min. The bees in the control groups ***(“Control”*)** in experiments 1, 2 & 3 were handled and chilled on ice (20-25 minutes) similarly to bees from the other experimental groups, but not treated with a drug or vehicle. **Sham-treated bees *(“Sham”*)** were similarly chilled and then treated on successive days with the castor oil and dimethylformamide (DMF) vehicles: On day 1 post-emergence, we applied the castor oil with an amount (4.0-5.2µl/bee) corrected for body size as detailed in Supplementary Table S1. On Day 2, in parallel to the JH-III treatments (see below), we treated the Sham treatment bees with DMF (3.5µl/ bee, irrespective of body size). All other treatments and handling conditions were kept similar to the other treatment groups. **Precocene-I treatment *(“P-I”*):** Given that the CA is the only known source of JH in insects (Riddiford, 2008), and that Precocnes cause atrophy of the corpora allata glands (Haunerland and Bowers, 1985), we used Precocene-I (P-I; Sigma-Aldrich, cat # 195855) to reduce circulating JH titers (for details see Shpigler et al. 2016). We suspended the P-I in castor oil (Sigma-Aldrich, cat # 259853) and applied the solution to the dorsal part of the thorax. The P-I + castor oil mixture, which is highly viscous, was thoroughly mixed with a pipette and vortexed at high speed for 2 minutes to better disperse the drug. The precise P-I dose was selected based on our previous studies (Shpigler et al., 2016) and further ad hoc validations for each new P-I batch that we used. These validations helped to assure a high survival rate and effective manipulation of JH levels (Fig.1A). Based on these considerations, we used 200-260µg per bee which was delivered in 4.0-5.2µl vehicle volume (Table S1). The volume was adjusted according to the bee body size such that the drug concentration remains similar at 50µg/µl/bee to minimize density-dependent adsorption of the drug. As an index for body size, we used the length of the front wing marginal cell (Shpigler et al., 2013; Yerushalmi, 2006). The cold-anesthetized and treated bees were placed on ice-chilled glass plates immobilized for ∼10 minutes after P-I treatments to allow the drugs to be absorbed, and minimize the incidents of wiping-off, as this can lead to variation in treatment affectivity. **Replacement therapy (*“P-I+JH-III”*):** we applied JH-III (Sigma-Aldrich, cat # J2000) dissolved in 3.5µl of Dimethylformamide (DMF, J.T Backers, cat # 7032) to bees that were already treated with P-I on the day before. This one day gap between the treatments was important for stress relief and efficacy of the two tandem treatments. We treated each bee with an amount of 50µg/bee JH-III, which we selected based on preliminary dose-response JH-III application studies and assessments of ovarian state (Fig.1B). This dose was as effective as higher doses used in previous studies with bumble bees in which 70 or 100µg/bee were applied on the thorax (Amsalem et al., 2014; Shpigler et al., 2014, 2016). P-I treated bees were collected and anesthetized by chilling on ice for 5-10 minutes. When the bee was motionless, we topically applied JH-III dissolved in DMF (replacement therapy). The treated bees were left immobilized on ice for additional ∼10 min for efficient penetration and for minimizing drug wiping off.

**Figure 1:**
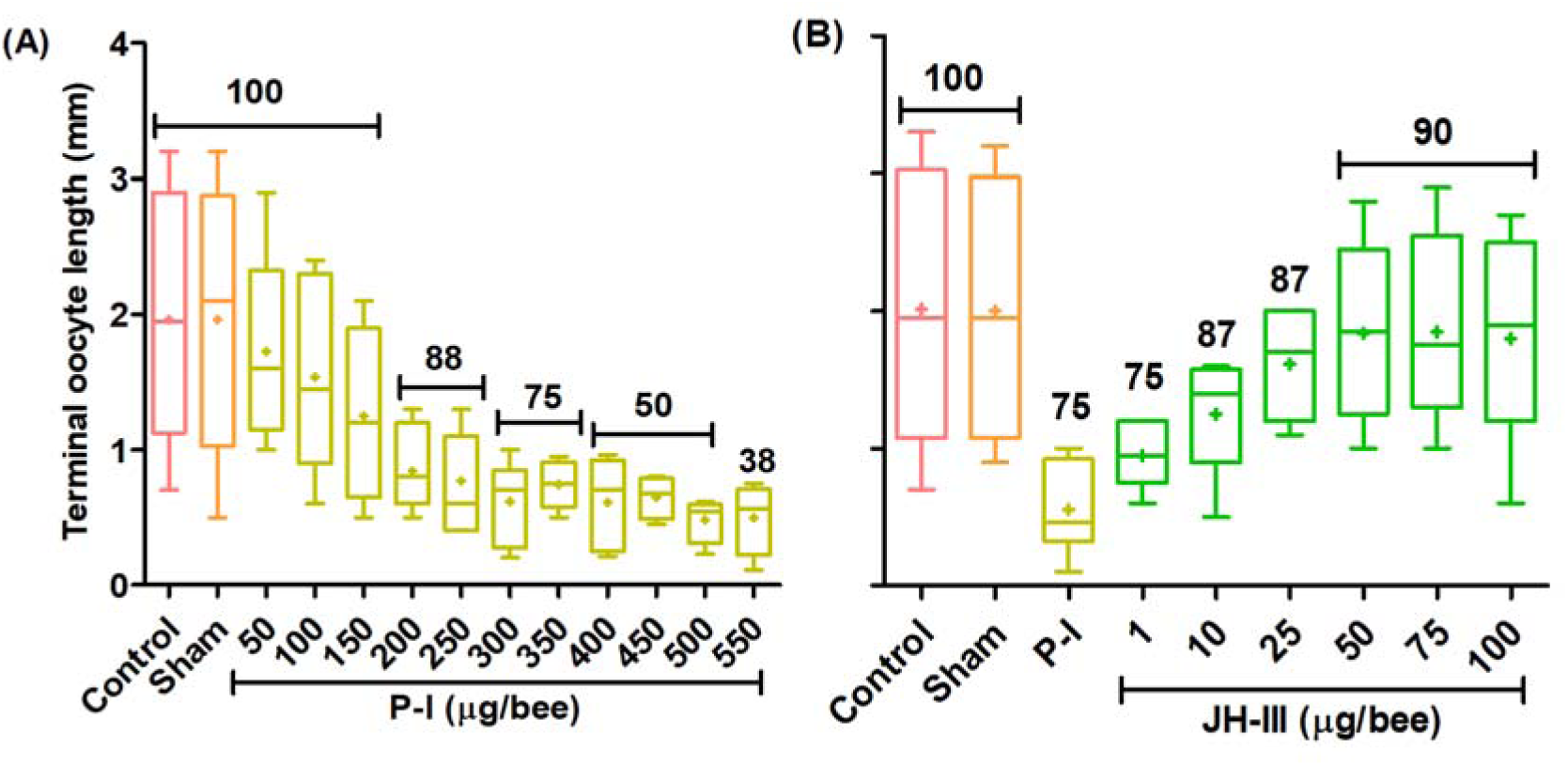
Dose-dependent effect of topical treatments with Precocene-I and Juvenile hormone on worker ovarian development: (A) Treatment with Precocene-I (P-I). The allatotoxin P-I was mixed with castor oil and amounts were adjusted according to the bee body size (4.0-5.2µl/ bee; see Table S1 for details). (**B)** Replacement therapy with juvenile hormone III (JH-III), the natural JH of bumble bees. In this experiment bees were first treated with P-I and then on the following day, with increasing amounts of JH-III (1.0-100µg/bee/3.5µl DMF). The sample size was 2-3 groups, each consisting of 4 worker bees (n=8-12 bees/ treatment). The box plots show the median (line) and mean (+), the box frame spans over the first to the third quartile. The whiskers depict the 5th/95th percentile, outliers are depicted with dots. The number above the bars show the percentage survival.

### 2.3. Behavioral observations

Focal orphan groups were observed twice a day during days 3-5 after the first treatment (a total of 120 min observations per group). Each observation day included a morning observation session between 09:00–11:00h, and an evening observation session between 16:00–18:00h. Each session lasted 20 minutes. Orphan (“*queenless*”) *B. terrestris* workers at this age typically show a high level of agonistic behaviors and establish clear dominance hierarchies (Amsalem and Hefetz, 2010; Bloch et al., 1996; Geva et al., 2005; Van Doorn, 1989). Given the importance of agonistic interactions for both the establishment and maintenance of dominance hierarchies in bumble bees, we recorded threatening displays that include *“buzzing”*, and *“pumping”* as described in Bloch et al., (1996). Buzzing displays are characterized by fast, short wing vibrations of a worker facing another bee; “pumping” is performed by a bee standing and facing a nestmate bee while performing distinct dorso-ventral pumping movement with her abdomen (Bloch et al., 1996; Duchateau, 1989; Geva et al., 2005). Dominant bees typically switched between buzzing and pumping behaviors. “Overt aggression”, which commonly follows threatening displays, includes darting and attacks directed towards another bee. To estimate the dominance of individual bees we used the Dominance Index (DI) that was calculated as in previous studies (Bloch et al., 1996; Geva et al., 2005; Van Doorn and Heringa, 1986). Briefly, the DI is the proportion of encounters between each pair of bees in which the focal bee did not retract, out of the total encounters [1-(Retractions/Total encounters)]. Thus, we used the amount of overt aggression and threatening displays as indices for aggressiveness, and the DI as an index for dominance. The dominance rank of a bee depicts its DI relative to that of others in the same queenless group. In each group, the worker with the highest DI value was termed “*alpha*” (*α*), following by “*beta*” (*β*), “*gamma*” (*γ*) and “*delta*” (*δ*) in descending DI order. Observers were blind to treatments. To ensure similar recording of focal behaviors across observers, all the observers were trained individually. The similarity in the behavioral recording was ensured by training sessions in which the instructor and trainee recorded behavioral observations in parallel. The observations were conducted under dim red light that bees do not see well. The observers entered the room carefully and avoided touching the tables on which the colonies were placed in order to avoid any movements or vibrations that can startle bumble bees. Observations were conducted silently and there were no visible signs that the bees were disturbed by the observations. The reported indices for each of these behaviors are the sum of total events recorded over 120 min of observations across 3 days (i.e. morning and evening observations on days 3, 4 and 5) and are presented as event per hour.

### 2.4. Assessment of ovarian state

At the end of each experiment, we stored the focal bees at −20^0^C. To assess ovarian state, we fixed the bee on a wax filled dissecting plate under a stereomicroscope (Nikon SMZ645) and immersed it in honey bee saline (Huang et al., 1991). We cut three incisions through the lateral and ventral abdominal cuticle using fine scissors and then immersed the internal organs in saline. We gently removed the ovaries into a drop of saline on a microscope slide, and use the ocular ruler to measure the length of the four largest terminal oocytes from both ovarioles. We used the average oocyte length of the four largest terminal oocytes as an index for ovarian state.

### 2.5. Experiment 1: The influence of JH on dominance and aggression in groups of similar size, differently-treated queenless worker bees

We collected callow bees (<24hrs of age) from various source colonies, determined their respective sizes, and assigned them randomly to one of the four manipulations described in Section 2.2. Following the treatments on days 1 & 2, we established groups on Day 3. Each orphan (“queenless”) group of four included one untreated (handling control) bee, one sham control bee, one P-I treated bee, and one replacement therapy treated bee. The four bees in the same group were selected to be of a similar body size. Bees were kept in isolation in a small Petri dish (35 × 12 mm) during the first and second day of the experiment (Section 2.2). The isolated bees were fed ad libitum with pollen cakes and sugar syrup and kept under controlled environmental conditions (29 ± 1 °C; 55% ± 10% RH). This procedure minimized the possible lateral transfer of treatment solutions (i.e., JH or P-I) across differentially treated groupmate bees. We made this adjustment in protocol following a preliminary experiment in which the ovaries of P-I treated bees were larger than expected in mixed groups that were constituted immediately after the first treatment (compare Fig. S1 and Fig. 2A). Each individual bee in this experiment was tagged with a different numbered colored disk allowing individual identification. The number tag (Opalith tags, Germany) was attached at the time of the second treatment. On Day 3 bees were assigned in groups of four per cage. Behavioral observations and oocyte measurements were performed as described in sections 2.3 & 2.4.

**Figure 2:**
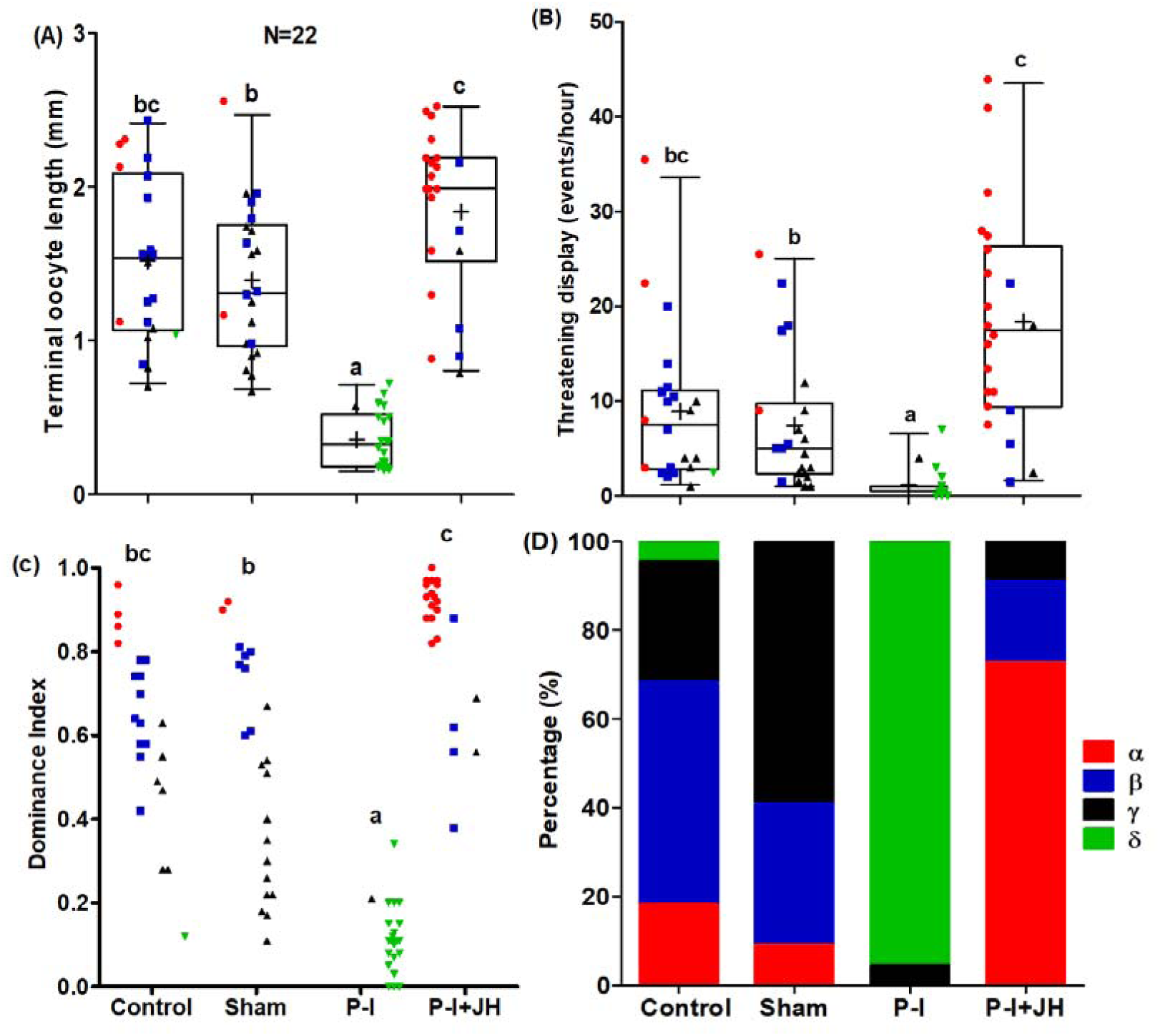
JH affects ovarian development, aggression and dominance status in groups of four orphan workers of similar body size and mixed treatments. (A) Ovarian state; (B) threatening displays; (C) Dominance Index. In A – C: Each colored symbol depicts the value of an individual worker bee. The color and shape of the symbols correspond to the, bee’s dominance rank within its group with α, β, γ & δ describing descending dominance ranks. Data were [Ln(X+1)] transformed to normalize the distribution for statistical analyses. Treatments marked with different small letters are statistically different in a Wilk’s Lambda test followed by Bonferroni’s correction. N=22 bees treated and tested per treatment; (D) Dominance rank distribution. The number in each stack box depicts the percentage of bees of each dominance rank. Treatment had a significant effect on the propensity to acquire a certain dominance rank (Chi-square test, p<0.001). Other details as in Fig.1.

### 2.6. Experiment 2: The influence of JH on dominance and aggression in groups of different size, similarly-treated queenless worker bees

We treated callow bees and assigned them to queenless groups within 1hr of collection from the source colony. Worker bees in each group differed in body size – each bee was chosen from a different size class (as denoted in Table S1; Two way ANOVA, F _(Size range) 3, 76_=546.8, p<0.05; F _(Treatment groups) 3, 76_=0.54, p>0.05, partial η*2*=0.99). The experimental outline was different in several ways from Experiment 1: (1) All the bees within a treatment group were treated similarly; (2) groups were constituted of bees having four different body sizes (Table S1). Each focal bee was paint marked with a different color to allow individual identification. The individual paint marks were applied to the dorsal part of the thorax soon after applying JH-III to the bee abdomen on Day 2. At this day, the castor oil on the thorax was already absorbed. After drying up, the bees of all the groups were reassembled into their respective wooden cages. Behavioral observations and oocyte measurements were performed as described in sections 2.3 & 2.4. This experimental design allowed us to test the effects of both body size and JH manipulation on ovarian development and behavior.

### 2.7. Experiment 3: The influence of JH on dominance and aggression in groups of similar size, similarly-treated queenless worker bees

The experimental outline is similar to Exp. 2 other than that the bees in each group were of similar body size (one way ANOVA, F_3, 100_=0.40, p>0.05, partial η*2*=0.01). This experiment allows us to test the effect of JH on aggression and dominance while controlling for the possibly confounding factors of size and age.

### 2.8. Experiment 4: The influence of JH on dominance and aggression in groups of same size bees in which dominance hierarchy has been already established

In contrast to the three previous experiments, here we treated bees that have already established a stable dominance hierarchy during the first four days following group constitution. We performed the first set of behavioral observations on days 3 & 4, before treating the bees. On the evening of Day 4, after determining the dominance rank within each group, we divided the groups into two sets. In one set, the α ranked individual was treated with P-I (500µg/5µl/ bee, henceforth termed “Experiment 4A”). In the other set of groups, we treated the δ rank individuals with JH-III (100µg/5µl/bee, henceforth termed “Experiment 4B”). We used here higher doses compared to the previous three experiments because the treated bees were older. Following treatment, we kept the bees on ice for 10 minutes for absorbing the treatment solution, followed with about 60 minutes of isolation in closed petri-dish with pollen and syrup to avoid the lateral transfer of the treatment solution. Each focal bee within a group was number tagged during group formation. The tag was attached to the dorsal part of the abdomen for the P-I treated bees, and on the thorax for JH-III treated bees. We performed additional sessions of behavioral observations on days 5 & 6 as described in section 2.3. Bees within the same group were of similar body size (One-way ANOVA, F_3, 132_= 0.06, p>0.05, partial η2 = 0.000). The bees were collected from several donor colonies and assigned randomly to treatment groups.

### 2.9. Statistical analyses

*Experiment 1*: We studied 22 Groups across three Trials with a total number of 88 bees. For each bee, we measured three variables: oocyte size, the number of threatening displays she performed, and their calculated dominance Index. For the statistical analyses, we used two-factorial General Linear Model, considering the Groups as independent units of sampling: One factor being the Trial, and another factor is the Treatment, having 4 repeated measures (Control, Sham, P-I, and P-I+JH). We thus have 3 × 4 = 12 random variables, and normality was ascertained for each after subjecting them to the ln(*x*+1) transformation.

*Experiment 2:* Here we studied 23 groups (5 Control, 5 Sham, 8 P-I, and 5 P-I+JH) with a total of 92 bees. The Groups are nested within the trials, which in turn are nested within Treatment. We analyzed three variables: oocyte length, the number of threatening displays, and the Dominance Index. Each was considered as the dependent variable in a two-way nested ANOVA. Normality was ascertained after subjecting each dependent variable to the square-root transformation. We also tested the effect of Treatment on the within-group dominance hierarchy, measured by the Dominance Range (i.e., the difference in Dominance Indices between α and δ bees in the group), using a two-way nested ANOVA.

*Experiment 3:* We studied 26 Groups (5 Control, 5 Sham, 8 P-I, and 8 P-I+JH) consisted of 104 bees. The statistical analysis is similar to that described for Experiment 2.

*Experiment 4:* In Experiment 4A we had 20 groups, 10 in which the α bee was treated with PI, and 10 in which the α bee was treated with only the vehicle (altogether 80 bees). In Experiment 4B we had 16 groups, 8 in which the δ bee was treated with JH-III, and 8 in which the δ bee was treated with only the vehicle (altogether 64 bees). For each bee we measured the terminal oocyte size, the number of threatening displays, and the DI at the first and at the second sets of behavioral observations. After applying the square-root transformation (for achieving normality), we considered for each bee three variables: (1) the size of terminal oocyte, (2) the difference between the number of threatening displays during the second and the first sets of behavioral observations, and (3) the difference between the dominance rank at the second and the first sets of behavioral observations.

For each dominance rank, we then compared in Experiment 4A the means of these three variables between the bees of the treated (P-I) and Control (vehicle) groups. Similarly, in Experiment 4B we compared for each dominance rank, the means of the abovementioned three variables between the bees of the treated (JH-III) and Control (vehicle) groups (all comparisons were performed using independent samples *t*-tests, assuming equal or unequal variances, as determined by the corresponding *F*-tests).

Dominance hierarchy as a function of JH level: We used three measures for assessing the linearity of the dominance hierarchy within a group: (1) The DI range, (2) the DI standard deviation, and (3) the inverse of the Shannon’s Diversity Index, which we calculated using the DI data. We used the ovarian state as a proxy for JH level (see above) and considered two measures: (1) The mean length of the four largest oocytes and (2) the length of only the largest oocyte in the ovaries. For testing the relationship between the extent of the dominance hierarchy and ovarian state, we performed six linear regressions, one for each of the 3 × 2 possible combinations. These were done twice – once on the data of Experiment 2 (different size bees) and once on the data of Experiment 3 (same size bees). We used partial η2 for effect size in ANOVA, whereas Cohen’s *d (Cd)* measure of effect size was calculated for pairwise comparisons.

## 3. Results

### 3.1. Experiment 1: The influence of JH on dominance and aggression in groups of similar size, differently-treated queenless worker bees

The assessment of ovarian state confirm that our JH manipulations were effective; P-I treated bees had ovaries containing smaller oocytes compared to control bees, an effect that was reverted by replacement therapy (Fig. 2A; Wilks’ Lambda = 0.04, *F*_3, 17_ = 130.45, *P* < 0.001, partial η^2^ = 0.96; Bonferroni adjusted pairwise comparisons, Control vs. Sham: *P* > 0.999, *Cd*=0.31; Control vs. PI: *P* < 0.001, *Cd*=3.76; Control vs. JH: *P* = 0.180, *Cd*=0.71; Sham vs. PI: *P* < 0.001, *Cd*=4.41; Sham vs. JH: *P* = 0.006, *Cd*=1.16; PI vs. JH: *P* < 0.001, *Cd*=4.64). Trial had no significant effect on oocyte size (*F*_2, 19_ = 0.54, *P* = 0.59, partial η^2^ =0.01). The P-I treated bees also showed reduced amounts of threatening displays, an effect that was recovered by the replacement therapy, showing that it is regulated by JH (Fig. 2B; Wilks’ Lambda = 0.11, *F*_3,17_ = 45.46, *P* < 0.001, partial η^2^ = 0.89; Bonferroni adjusted pairwise comparisons, Control vs. Sham: *P* > 0.999, *Cd*=0.38; Control vs. PI: *P* < 0.001, *Cd*=2.46; Control vs. JH: *P* = 0.061, *Cd*=0.86; Sham vs. PI: *P* < 0.001, *Cd*=2.37; Sham vs. JH: *P* < 0.001, *Cd*=1.54; PI vs. JH: *P* < 0.001, *Cd*=3.51). There was a small but statistically significant effect of Trial (*F*_2, 19_ = 6.68, *P* = 0.006, partial η^2^ =0.07). JH manipulation also had a significant influence on the Dominance Index (Fig. 2C; Wilk’s Lambda = 0.03, *F*_3, 17_ = 166.90, *P* < 0.001, partial η^2^ = 0.96; Bonferroni adjusted pairwise comparisons, Control vs. Sham: *P* = 0.944, *Cd*=0.44; Control vs. PI: *P* < 0.001, *Cd*=3.43; Control vs. JH: *P* = 0.054, *Cd=0.88*; Sham vs. PI: *P* < 0.001, *Cd* =2.12; Sham vs. JH: *P* = 0.003, *Cd*=1.26; PI vs. JH: *P* < 0.001, *Cd*=5.75) with no effect of Trial *(F*_2, 19_ = 2.78, *P* = 0.087, partial η^2^=0.01). The Bonferroni adjusted pairwise comparisons (Fig. 2C) show that the P-I treated bees had a significantly lower Dominance Index. Thus, the rank in the dominance hierarchy was influenced by treatment (Chi-Square Test _(n=22)_ =114.55, df=9, p<0.001). In all groups except one, the P-I treated bee showed the lowest dominance rank (δ; 95.45%), and in the remaining one group it had the second lowest (γ) rank. The majority of the replacement therapy treated bees were the most dominant in their group (73%; Fig. 2D).

The replacement therapy not only recovered the P-I treatment effect, but actually resulted in better-developed ovaries and higher levels of threatening displays and dominance index compared to the sham-treated (but not the control bees, Fig. 2). This experiment shows that JH can influence the amount of threatening displays and dominance rank in groups composed of workers bees with similar body size.

### 3.2. Experiment 2: The influence of JH on dominance and aggression in groups of different size, similarly-treated queenless worker bees

Ovarian state differed between treatment groups, confirming the affectivity of JH manipulations in this experiment (Fig.3A, Treatment: *F*_3, 69_ = 21.91, *P* < 0.001, partial η^2^ = 0.49; Group: *F*_19, 69_ = 0.71, *P* = 0.793, partial η^2^ = 0.16; Bonferroni adjusted pairwise comparisons, Control vs. Sham: *P* > 0.999, *Cd*=0.12; Control vs. PI: *P* < 0.001, *Cd*=1.69; Control vs. PI+JH: *P* > 0.999, *Cd*=0.11; Sham vs. PI: *P* < 0.001, *Cd*=1.82; Sham vs. PI+JH: *P* > 0.999, *Cd*=0.01; PI vs. PI+JH: *P* < 0.001, *Cd*=1.81). All the bees in the P-I treatment groups, including the most dominant individuals, had undeveloped ovaries, similar in size to those of the low ranked bees (γ & δ) in the Control and Sham treatment groups (Fig. 3A). In the Replacement Therapy groups, all the bees, including the low ranked individuals, had similarly active ovaries (One way ANOVA _(replacement group)_, F_3,16_=1.17, p=0.35, partial η*2*=0.18). The average oocyte length of the γ and δ ranked individuals in the replacement therapy groups appeared larger than for bees of similar dominance rank in the Control, Sham and P-I treatment groups (Fig. 3A).

**Figure 3.**
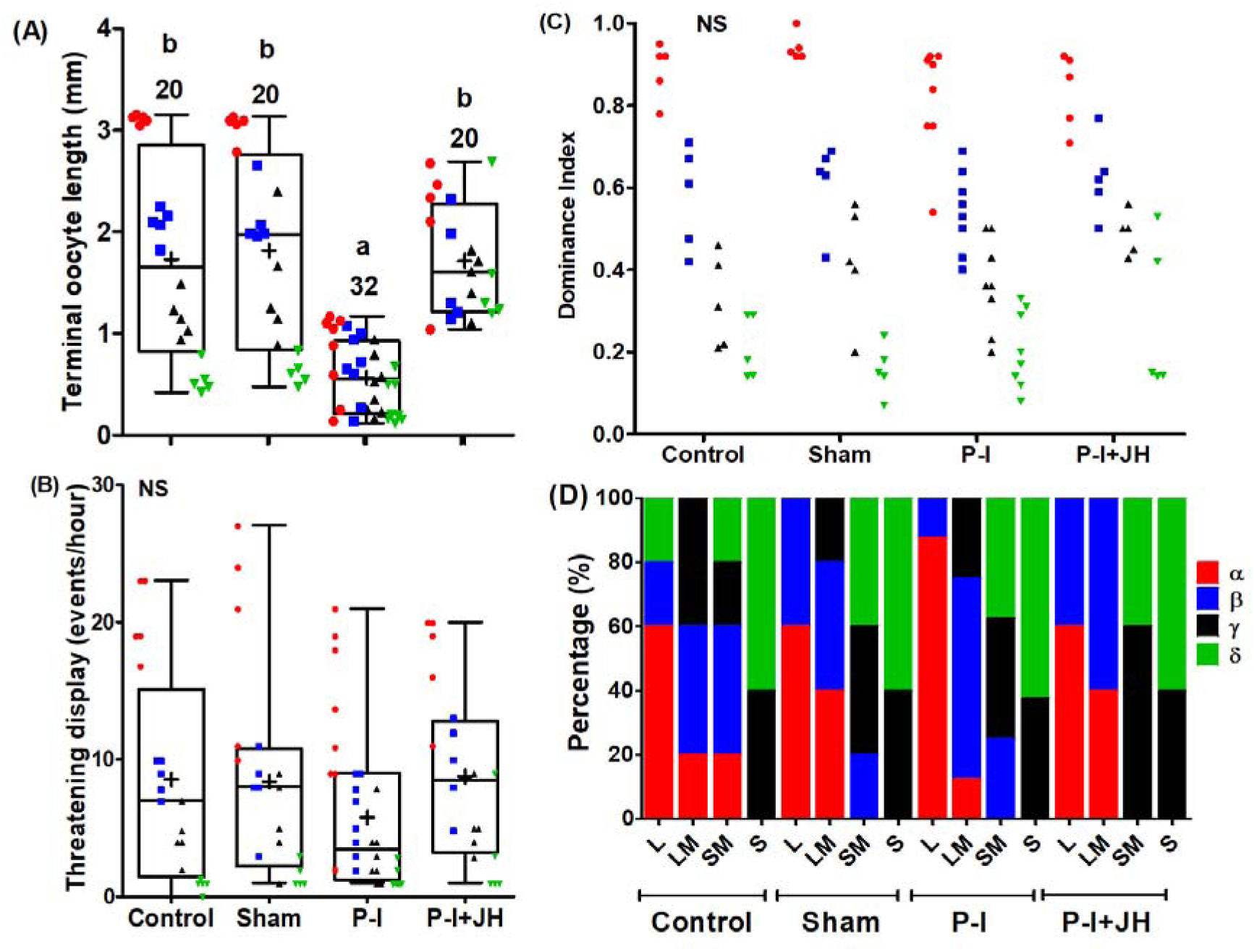
JH effects on ovarian development, dominance status and threatening displays in groups of four orphan workers of different body size and similar treatment. (A) Ovarian state (B) Threatening behavior, (C) Dominant Index, and (D) Dominance rank distribution. Data were square-root transformed to normalize the distribution for statistical analysis. Letters (L, LM, SM, and S) refer to the body size of the bees (Table S1). Other details as in Fig. 1 & 2.

Threatening displays did not differ between treatment groups (Fig. 3B, *F*_3, 69_ = 0.99, *P* = 0.403, partial η^2^ = 0.04) and there was no significant effect for group factor (Fig. 3B, *F*_19, 69_ = 0.57, *P* = 0.916, partial η^2^ = 0.14). In all groups, the most dominant individual performed most of the threatening displays (Fig. 3B). All the treatment groups established clear dominance hierarchy (Fig. 3C; Two way Nested ANOVA _(Treatment)_, F_3, 69_=0.30, p > 0.05, partial η*2*=0.01). There was also no significant effect of treatment on the Dominance Index (Fig.3C, *F*_3, 69_ = 0.30, *P* = 0.823, partial η^2^ = 0.01) and no significant effect of the Group factor (Fig.3C, *F*_19, 69_ = 0.16, *P* > 0.999, partial η^2^ = 0.04).

To compare the strength of the linear dominance hierarchies across treatments, we first estimated the range of dominance indexes by subtracting the dominance index of δ from that of the α individual in each group. This value is expected to be larger when the dominance hierarchy is clear. We found no significant effect of treatment on Dominance Range (Fig.S3, *F*_3,12_ = 1.02, *P* = 0.418, partial η^2^ =0.20) and no effect of the Trial (*F*_7, 12_ = 1.96, *P* = 0.147, partial η^2^ = 0.53). We next tested the correlation between ovary state (length of the largest oocyte, or mean of the four largest oocyte) as a proxy for JH titer and three different indices for the strength of the dominance hierarchy: The range of the Dominance Index (see above), the Standard Deviation of the Dominance Index and the inverse of the Shannon’s Diversity Index. We found that none of the six combinations produced a statistically significant correlation between the measure of the dominance hierarchy and that of ovarian state (Table S2).

Taken together, these analyses show that bees subjected to similar JH manipulation and thus, have similar ovarian state, can nevertheless establish clear dominance hierarchies, similar to that observed in groups of workers in which JH levels were not manipulated (i.e., the Control and Sham groups). Dominance rank in all treatment groups was strongly influenced by body size with the largest individual most likely to acquire the top tank in the dominance hierarchy, and the smaller individuals typically occupying the lower ranks. Importantly, these include the P-I and replacement therapy treatment in which all the bees in a group are assumed to have overall similar JH titers. In 7 out of 8 P-I treated groups (87.5%, Fig. 3D, Chi-Square Test _(***P-I***)_ =448.5, df=9, p<0.001) and in 3 out of 5 replacement therapy treated groups the largest individual was the most dominant in the group (60%, Fig. 3D, Chi-Square Test _(***P-I+JH***)_ =432.00, df=9, p<0.001). This proportion is comparable to the Control and sham groups (3 out of 5 groups = 60%, Fig. 3D, Chi-Square Test _(***Control***)_ =240.00, df=9, p<0.001, Chi-Square Test _(***Sham***)_ =304.00, df=9, p<0.001). This experiment shows that body size has a strong influence on agonistic behavior and dominance rank in small groups of queenless workers, and this effect of body size is not compromised in groups in which all bees are manipulated to have similar JH titers and level of ovarian activation.

### 3.3. Experiment 3: The influence of JH on dominance and aggression in groups of similar size, similarly-treated orphan worker bees

Given that Exp.1 and Exp.2 indicate that both JH and body size (respectively) influence dominance rank in small queenless groups, we next studied dominance hierarchy formation in groups of bees of similar body size and JH levels. The strong influence of treatment on ovarian state confirmed the effectiveness of our JH manipulations (Fig. 4A; Treatment: *F*_3, 78_ = 43.78, *P* < 0.001, partial η^2^ = 0.63; Group: *F*_22,78_ = 0.35, *P* = 0.996, partial η^2^ = 0.09; Bonferroni adjusted pairwise comparisons, Control vs. Sham: *P* > 0.999, *Cd*=0.15; Control vs. PI: *P* < 0.001, *Cd*=2.07; Control vs. PI+JH: *P* = 0.114, *Cd*=0.68; Sham vs. PI: *P* < 0.001, *Cd*=1.93; Sham vs. PI+JH: *P* = 0.028, *Cd*=0.83; PI vs. PI+JH: *P* < 0.001, *Cd*=2.76). Even the most dominant P-I treated individuals had undeveloped ovaries, with the largest oocytes appearing similar in size to those of subordinate bees in the Control and Sham treatment groups. Similarly, all the bees in the Replacement Therapy treatment groups, including those with low dominance rank, had active ovaries appearing similar to dominant bees in the Control and Sham treatment groups (Fig. 4A). These ovarian analyses suggest that in both the P-I and Replacement Therapy groups all four individuals had comparable, either low or high, JH levels, respectively (Fig.4A; One way ANOVA within groups: P-I -- F=1.70, df=3, p>0.05, partial η*2*=0.15; Replacement Therapy -- F=29.90, df=3, p<0.0001, partial η2=0.76).

**Figure 4:**
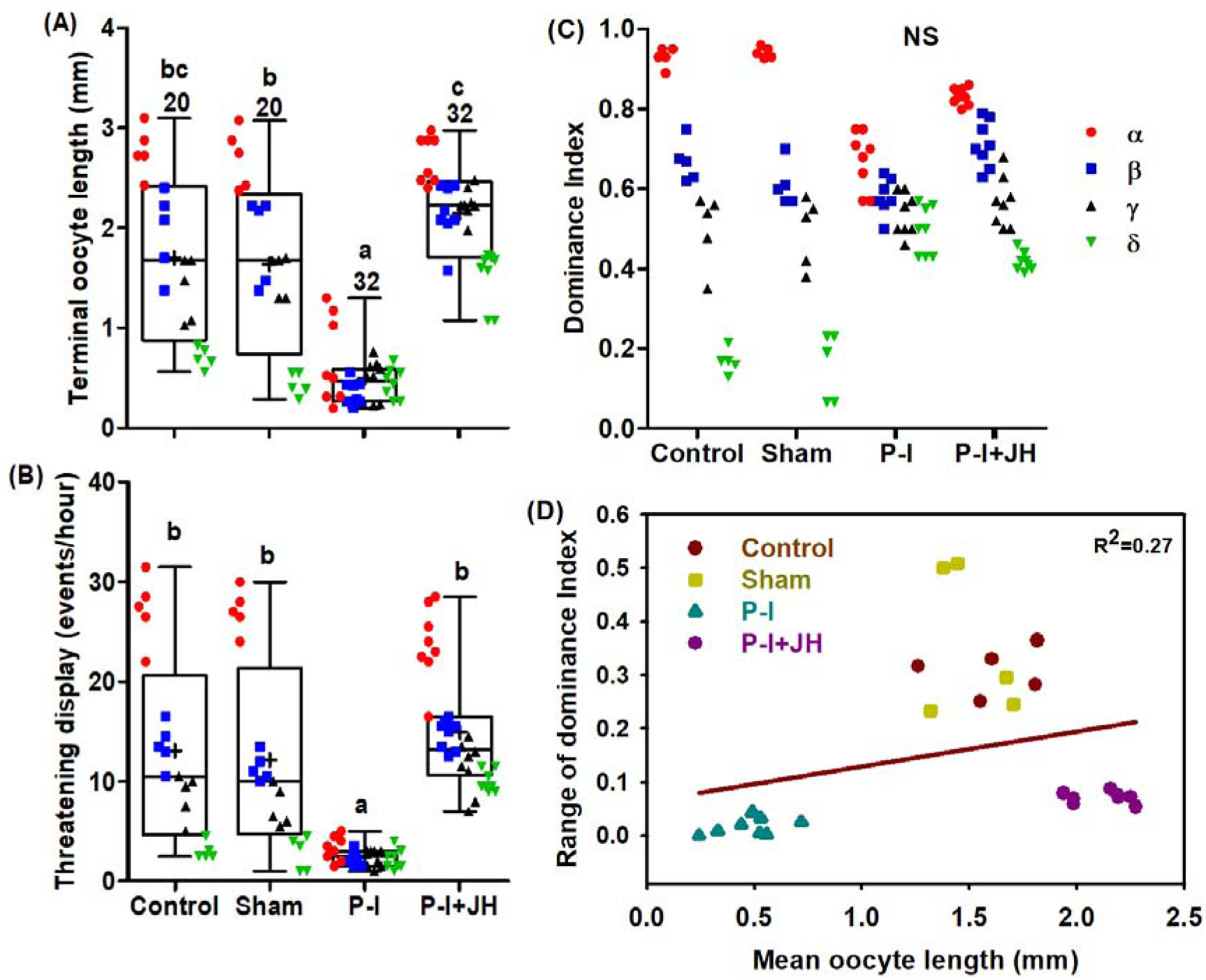
JH effects on ovarian development, dominance rank and threatening displays in groups of four orphan workers of similar body size and treatment. (A) ovarian state, (B) threatening displays, (C) Dominance Index and (D) Correlation between mean oocyte length and range of dominance index. Data were square-root transformed to normalize the distribution for statistical analysis. Other details as in Fig. 1&2.

JH manipulation also affected threatening behavior. P-I treated bees performed overall less, and replacement therapy treated bees more, threatening displays compared to the Sham and Control groups (Fig. 4B, Treatment: *F*_3, 78_ = 26.81, *P* < 0.001, partial η^2^ = 0.51; Group: *F*_22, 78_ = 0.10, *P* > 0.999, partial η^2^ = 0.03; Bonferroni adjusted pairwise comparisons, Control vs. Sham: *P* > 0.999, *Cd*=0.19; Control vs. PI: *P* < 0.001, *Cd*=1.73; Control vs. PI+JH: *P* > 0.999, *Cd*=0.38; Sham vs. PI: *P* < 0.001, *Cd*=1.55; Sham vs. PI+JH: *P* = 0.301, *Cd*=0.57; PI vs. PI+JH: *P* < 0.001, *Cd*=2.12). These effects of JH treatment were closely linked to dominance rank: The frequency of threatening displays was severely reduced in high-rank individuals in the P-I treated groups (Fig. 4B) and elevated in low ranked individuals in the Replacement Therapy treatment groups. JH manipulation did not affect Dominance Index (Fig. 4C; *F*_3, 78_ = 0.67, *P* = 0.574, partial η^2^ = 0.03; Group effect - *F*_22, 78_ = 0.07, *P* > 0.999, partial η^2^ = 0.02). It is remarkable that despite their similar JH levels, ovary state, and elevated threatening displays by subordinate individuals, bees in the Replacement Therapy groups nevertheless differ in their Dominance Index (Fig. 4C; One way ANOVA within group, F=122.77, df=3, p<0.001, partial η*2*=0.93). Although the variation was lower in groups of P-I treated bees, the Dominance Index was nevertheless significantly different among bees differing in their dominance rank (Fig. 4C; F_3, 28_=13.41, p<0.001, partial η^2^ = 0.50). The Dominance Range was influenced by treatment (*F*_3, 14_ = 150.82, *P* < 0.001, partial η^2^ = 0.97; Trial effect − *F*_8, 14_ = 2.38, *P* = 0.074, partial η^2^ = 0.58), and was significantly lower in the P-I and Replacement Therapy groups compared with the Control and the Sham groups (Fig. 4C, D; Bonferroni adjusted pairwise comparisons, Control vs. Sham: *P* > 0.999, *Cd*=0.38; Control vs. PI: *P* < 0.001, *Cd*=10.02; Control vs. PI+JH: *P* < 0.001, *Cd*=5.94; Sham vs. PI: *P* < 0.001, *Cd*=10.39; Sham vs. PI+JH: *P* < 0.001, *Cd*=6.32; PI vs. PI+JH: *P* < 0.001, *Cd*=4.07). By contrast to Exp. 2, here the strength of the dominance hierarchy was positively correlated with the ovarian state when estimated using the range of dominance index, Shannon’s diversity index, or the standard deviation of dominance index (Fig. 4D; Table S3). These results suggest that even in groups composed of workers with similar body size, age, and JH levels, there is a weak dominance hierarchy.

### 3.4. Experiment 4: The influence of JH on dominance and aggression in groups of same size bees in which dominance hierarchy has been already established

Here, we tested whether P-I treatment can decrease the rank of dominant individuals (Exp. 4A), and whether JH treatment can increase, the dominance of subordinate (Exp. 4B), in groups that had already established stable dominance hierarchies. Dominant (α) individuals that were treated with P-I had less developed ovaries compared to same age dominant bees that were treated with only the vehicle (Fig. 5A, left; Table 1Ai). It is notable, however, that the P-I treated dominant bees still have developed ovaries when assessed at 7 days of age, suggesting that reducing JH levels at this stage only slows or prevents further growth, but does not lead to a significant decrease in oocyte size. The ovarian state was similar for the β, γ & δ ranked bees (that were not treated) in the Sham and P-I treatment groups (Fig. 5A, left; Table 1Ai). The P-I treated dominant bees did not show a decrease in the amount of threatening displays (Fig. 5B, left; Table 1Aii) or DI (Fig.5C, left; Table 1Aiii). None of the α-ranked individuals lost their high dominance rank after the P-I treatment. Treating α individual with P-I also did not affect the level of threatening displays or the Dominance Index of the other bees in the group.

**Table 1A.** The influence of treating the α ranked individual with P-I on (i) the terminal oocyte length, (ii) the *change* in the number of threatening displays and (iii) the *change* in the dominance index for all bees in the group as function of their dominance rank (variables were square-root transformed for the statistical analyses).

| Dominance Rank | $\alpha$ | B | $\gamma$ | $\delta$ |
| --- | --- | --- | --- | --- |
| (i). Terminal Oocyte Length | $t_{11} = -4.66$<br>$P < 0.001$<br>$Cd=2.08$ | $t_{12} = 0.80$<br>$P = 0.440$<br>$Cd=0.36$ | $t_{14} = -0.52$<br>$P = 0.607$<br>$Cd=0.23$ | $t_{18} = -1.82$<br>$P = 0.086$<br>$Cd=0.81$ |
| (ii). Threatening Displays | $t_{12} = -0.69$<br>$P = 0.503$<br>$Cd=0.31$ | $t_{18} = 0.94$<br>$P = 0.359$<br>$Cd=0.42$ | $t_{18} = 1.62$<br>$P = 0.122$<br>$Cd=0.73$ | $t_{18} = -0.44$<br>$P = 0.663$<br>$Cd=0.20$ |
| (iii). Dominance Index | $t_{12} = -0.35$<br>$P = 0.729$<br>$Cd=0.16$ | $t_{10} = -1.28$<br>$P = 0.228$<br>$Cd=0.57$ | $t_{14} = 0.61$<br>$P = 0.548$<br>$Cd=0.27$ | $t_{10} = -0.73$<br>$P = 0.480$<br>$Cd=0.33$ |

\*Cd=Cohen's d

**Table 2B.** The influence of treating the δ ranked individual with JH-III on (i) the terminal oocyte length, (ii) the *change* in the number of threatening displays and (iii) the *change* in the dominance index for all bees in the group as function of their dominance rank (variables were square-root transformed for the statistical analyses).

| Dominance Rank | $\alpha$ | B | $\gamma$ | $\delta$ |
| --- | --- | --- | --- | --- |
| (i). Terminal Oocyte Length | $t_{14} = -1.44$<br>$P = 0.172$<br>$Cd=0.72$ | $t_{14} = -0.15$<br>$P = 0.883$<br>$Cd=0.08$ | $t_{14} = 1.57$<br>$P = 0.139$<br>$Cd=0.78$ | $t_{14} = 8.21$<br>$P = 10^{-6}$<br>$Cd=4.11$ |
| (ii). Threatening Displays | $t_{14} = 0.50$<br>$P = 0.629$<br>$Cd=0.25$ | $t_{14} = 1.36$<br>$P = 0.196$<br>$Cd=0.68$ | $t_{14} = -0.68$<br>$P = 0.508$<br>$Cd=0.34$ | $t_{14} = 5.25$<br>$P = 10^{-4}$<br>$Cd=2.62$ |
| (iii). Dominance Index | $t_{14} = 0.05$<br>$P = 0.958$<br>$Cd=0.03$ | $t_{14} = -1.12$<br>$P = 0.281$<br>$Cd=0.56$ | $t_{10} = 0.63$<br>$P = 0.543$<br>$Cd=0.31$ | $t_{14} = 3.98$<br>$P = 0.001$<br>$Cd=1.99$ |

\*cd=Cohen's d

**Figure 5:**
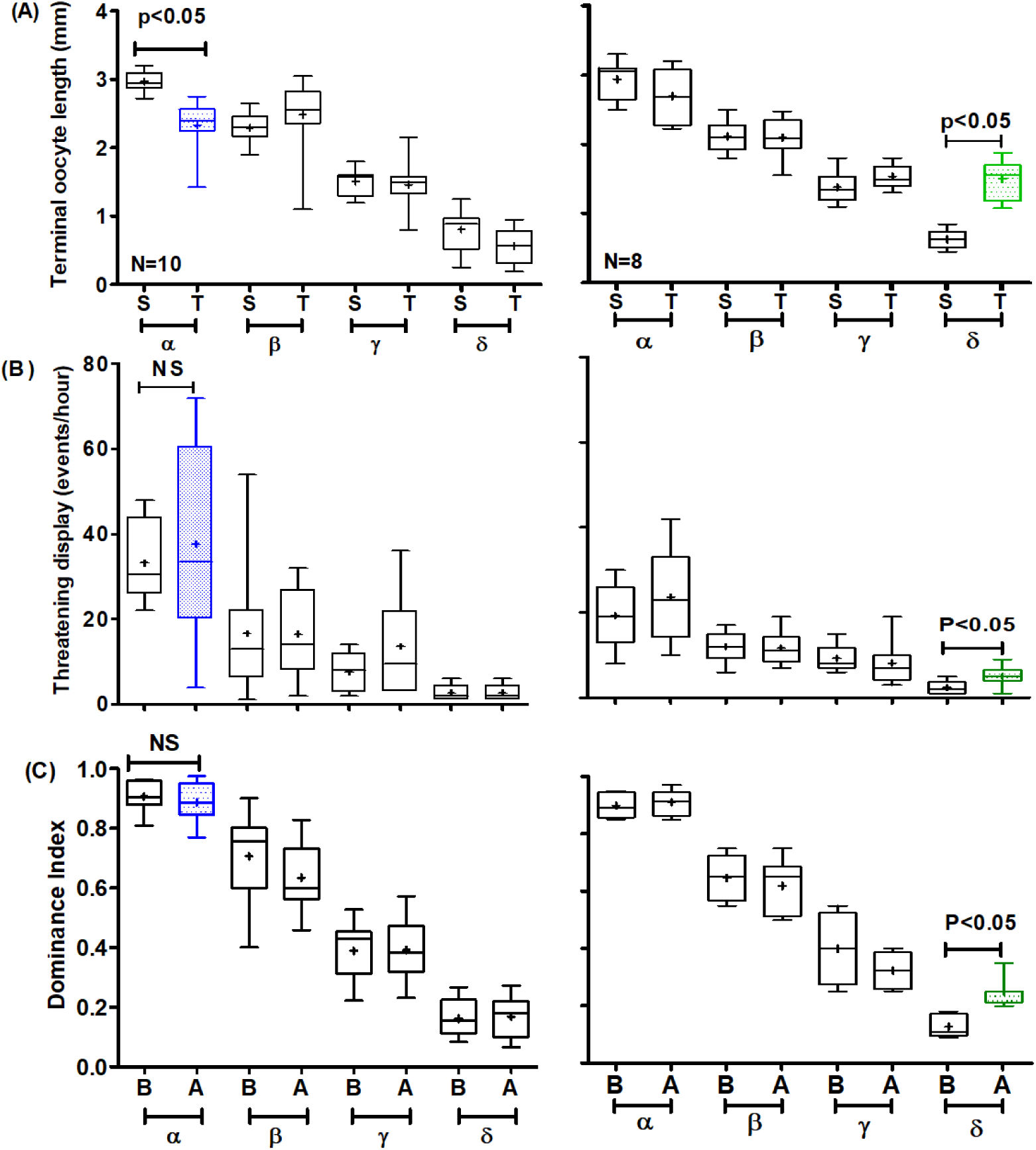
Effect of JH on dominance rank in groups that have already established dominance hierarchies. Left column – Exp. 4A in which we reduced JH levels for the most dominant individual; right column – Exp. 4B in which we elevated JH for the most subordinate individual. (**A**) ovarian state, (**B**) threatening displays. and (**C**) Dominance Index. The letters on the x-axis show the treatment groups: S=Sham, T=Treatment (P-I or JH), B=before the treatment and A=after the treatment). Data were square-root transformed to normalize the distribution for statistical analysis. Other details as in Figs. 1 and 2.

In Experiment 4B, we found that δ-ranked individuals that were treated with JH had significantly better-developed ovaries compared to sham-treated δ individuals in control groups suggesting that our manipulation successfully elevated circulating JH titers (Fig. 5A, right column; Table 1Bi). JH treatment also caused a significant increase in the frequency of threatening displays performed by the treated individual but not by her three groupmates (Fig. 5B, right column; Table 1Bii). This change in behavior was associated with a higher DI for the treated individuals when compared with the values before the treatment (Fig. 5C right column; Table 1Biii). Dominance rank comparison showed that in 1 out of the 8 (12.5%) groups the JH treated δ showed an increase in dominance rank, but this is not significantly different than the probability to change rank in the control groups in which the δ-ranked bee was treated with vehicle alone (0 out of the 8 groups, Fisher’s exact test _(δ -treated)_, p=0.54).

The correlation between the DI standard deviation before and after the treatment did not show any significant effect for α and δ treated individuals. This experiment suggests that JH has little, or only limited, effect on dominance rank after a stable dominance hierarchy had already been established.

## 4. Discussion

We combined treatments with the allatoxin P-I and replacement therapy with the natural JH to manipulate circulating JH levels in orphan groups of bumble bee workers. Our measurements of ovarian activity (terminal oocyte length) confirm that our JH manipulations were effective. Using this powerful system we show that in addition to its gonadotrophic effects on physiology, JH also affects social behaviors that are associated with reproduction such as aggressiveness and dominance. Our results suggest that JH interacts with additional factors such as body size and previous experience that also influence aggressiveness and dominance. These findings explain previous ambiguities concerning whether or not JH influences dominance in bumble bees.

In orphan quartets in which each callow worker was subjected to a different JH manipulation (Exp. 1), the P-I and replacement therapy treated bees typically show the lowest and highest dominance rank, respectively. These findings are consistent with the premise that JH influences dominance in this species. However, in groups in which callow bees of a similar size were subjected to the same treatment (Exp. 3), dominance rank was apparent even in the P-I and replacement therapy quartets in which all four bees were subjected to similar JH manipulation and had comparable ovarian activity, although dominance hierarchy was not as pronounced as in the Control and Sham treatment groups (Fig. 4C). These findings are consistent with the hypothesis that variation in JH levels is not necessary for the establishment of dominance hierarchies. This notion received further support in Exp. 2 in which the bees in the same group differ in body size, but all of which were subjected to the same treatment. Here, the dominance hierarchies in the P-I and replacement therapy treated groups were overall comparable to that in the Control and Sham treatment groups (Fig. 3C). Large body size increased and small size decreased, the propensity to secure the top dominance rank. These findings show that variation in body size is sufficient for establishing stable dominance hierarchies among bees that do not differ in JH titers and ovarian state. In the last experiment, we treated somewhat older (4-day-old) bees that have already established dominance hierarchies. We downregulated JH titers for the most dominant individual (α) and elevated JH titers for the most subordinate (γ) individual. These treatments affected ovarian state but had only limited effects on the level of threatening behavior and on the dominance rank of the treated individual. This experiment exemplifies the importance of previous experiment by showing that the influence of JH on dominance and aggression is small at best in bees that have already established a stable dominance hierarchy (Fig. 5). Taken together, our results show that JH is sufficient but not necessary for the regulation of aggressiveness, and the establishment and maintenance of a dominance hierarchy in *B. terrestris*. Body size, previous experience, and conceivably additional factors, also influence dominance status.

Our findings are consistent with the notion that JH modulates aggressiveness in bumble bees. In the first three experiments, bees treated with P-I to reduce JH titers showed less threatening displays relative to control and sham-treated bee’s (this trend was not statistically significant in Exp. 2), an effect that was reverted by replacement therapy with JH-III. JH manipulation however, did not affect aggressiveness in the fourth experiment, in which bees were treated after the dominance hierarchy had been already established. These manipulation experiments are consistent with the typical correlation between dominance rank and JH hemolymph titer (Bloch et al., 2000a; Bloch and Hefetz, 1999), in vitro JH biosynthesis rates (Bloch et al., 1996; Larrere and Couillaud, 1993), CA volume (Röseler et al., 1984; Van Doorn, 1989), and brain transcript abundance of the transcription factor *Kr-h1* which is considered a major JH readout gene in insects (Shpigler et al., 2010). Our findings are also consistent with studies showing that JH modulates aggression in other species of insects (Barth et al., 1975; Hunt, 2007; Kou et al., 2009, 2008; Pearce et al., 2001; Robinson et al., 1987; Röseler et al., 1984; Scott, 2006a, 2006b; Tibbetts and Huang, 2010), including other bees (Breed, 1983; Huang et al., 1994). It should be noted however, that some previous findings are not consistent with the hypothesis that JH affects dominance or aggression in bumble bees. Amsalem et al., (2014a, b) reported that oral treatment with JH, P-I, or both did not affect Vg mRNA levels and aggression, and suggested an alternative hypothesis stating that Vg regulates aggression independent of JH. Some of the inconsistencies between our results and that of Amsalem et al., (2014a, b), may be explained by methodological differences. For example, Amsalem et al., (2014a, b) used oral P-I treatment (3 mg/bee) and a higher JH-III dose (100 µg/bee compared to 50µg/bee in our study), that may affect the response to JH treatment (Pinto et al., 2000; Rutz et al., 1976). In addition, the analyses of ovarian activity in Amsalem et al., (2014b) suggest that their replacement therapy did not fully recover the effect of the P-I treatment in some of their experiments (e.g., Figs 2 and 3 in this paper). Our procedures also differ from that of Van Doorn (1989) who injected JH-I into one individual in a queenless group. He reported only a small effect on the propensity of the injected bee to acquire the top dominance rank. However, given that control bees were not disturbed in this experiment, it is possible that the stress of injection interfered with the hormonal influence; it is also not clear at what age the bees were injected. Similarly, JH was not found necessary for the expression of dominance behavior in the wasp *Synoeca surinama* (Kelstrup et al., 2014).

Our study points to factors other than JH that also affect dominance and aggression. Bees manipulated to have similar JH levels (i.e., all workers in the same group are similarly treated with P-I or replacement therapy) nevertheless established dominance hierarchies (Exp. 2 and 3). Exp. 2 further indicates that one important factor is body size. Bees in the P-I and Replacement Therapy groups showed agnostic behaviors and formed clear dominance hierarchy comparable to those in the control and sham-treated groups, despite having similarly developed ovaries (which is a proxy for JH titers; Fig. 3). Dominance rank in this experiment was not correlated with the ovarian state but rather with body size. These findings are consistent with earlier studies showing that body size affects dominance in untreated *B. terrestris* (Röseler, 1977; Van Doorn, 1989) and *B. atratus* worker bees (Matos and Garofalo, 1995). Body size is also correlated with dominance rank in other social bees (e.g., carpenter bees, Withee and Rehan, (2016), Lawson et al., (2017); halictine bee, Smith and Weller, (1989)) as well as other social insects such as paper wasps (Cervo et al., 2008; Chandrashekara and Gadagkar, 1991; Zanette and Field, 2009) and queenless ants (Heinze and Oberstadt, 1999; Nowbahari et al., 1999; Trible and Kronauer, 2017). The common correlation between body size and dominance rank can be explained by the importance of body size in determining the outcome of agonistic interactions and the establishment of dominance hierarchies. However, it should be noted that size is not always correlated with dominance rank in studies with bumble bees (Amsalem and Hefetz, 2010; Foster et al., 2004; Free, 1955) and other insects such as the paper wasp, *Polybia occidentalis* (O’Donnell and Jeanne, 1995). Size and aggression do not seem to be important for reproductive dominance in the primitively eusocial wasp *Ropalidia marginata* (Gadagkar, 2009; Bang and Gadagkar, 2012).

Previous experience can also influence the outcome of agonistic interactions and may explain the little effect of JH manipulation in Exp. 4. In many animals, older (but not too old) individuals are more likely to dominate younger opponents (Hughes and Strassmann, 1988; Khazraïe and Campan, 1999; Stevenson and Schildberger, 2013; Tsuji and Tsuji, 2005). More specific effects of previous agonistic interactions, known as “winner” and “loser” effects, reflect the increasing tendency of previous winners to win again, and of previous losers to be beaten in a conflict following a defeat (Goubault et al., 2019; reviewed in Hsu et al., 2006; Kim et al., 2018), and were recently suggested to be also important in a social wasp (Bang and Gadagkar, 2016). In most groups in Experiment 4, reducing JH and ovarian activity in the α-ranked bee did not cause her to lose her top position. Given α that the bees in each group were of a similar body size, it seems unlikely that the α bee kept her top rank after JH was reduced, and the δ bee remained the most subordinate after her JH titers were increased because they were the largest or smallest in their groups, respectively. We suggest that a more likely explanation is that previous experience as the dominant or subordinate individual, or both, have a stronger influence on subsequent dominance rank than our manipulation of circulating JH titers. The prior experience was found to affect dominance rank in many animals, including species of insects such as crickets and beetles (Arneson and Wcislo, 2003; Goubault and Decuignière, 2012; Khazraïe and Campan, 1999; Lee et al., 2014). To the best of our knowledge, loser and winner effects have not been explicitly tested in bumble bees, but there is evidence lending credence to this notion. For example, in queenless groups composed of both callow and older dominant workers, the later ones typically attain the top rank and inhibit ovarian development in the callows, but in groups composed of only callows, one of the youngsters acquires the top dominance rank (Bloch and Hefetz, 1999).

Our finding that JH is one of several factors affecting dominance and aggression in bumble bees can explain the apparently conflicting findings in previous studies (Amsalem et al., 2014b, 2014a; Röseler, 1977; Van Doorn, 1989). The effects of JH manipulation is influenced by the context, including JH titers of other bees in the group. This multiplicity of factors raises questions concerning the ways these factors interact during dominance hierarchy formation. We suggest that in groups of callow workers, hemolymph JH titers are less important during the initial stages of hierarchy formation. This premise is consistent with observations that the highest aggression and the initial establishment of a clear dominance hierarchy in small groups of callow workers typically occur during the first few days in which both JH biosynthesis rates (as determined in vitro) and JH titers are low (Bloch et al., 2000a, 1996; Van Doorn, 1989). Body size, aggressiveness, age, and previous experience may all play a role during this initial stage. We speculate that the burst of intensive agonistic interactions leads to the activation of the CA in the more aggressive and dominant individuals. This idea is consistent with the “challenge hypothesis”, which was developed to explain the social modulation of testosterone to increase aggression in the context of courtship and reproduction in vertebrates (Wingfield et al., 1990), and later extended to JH in insects (Scott, 2006a; Tibbetts and Huang, 2010; Bernstein et al., 1979; Kou et al., 2019, 2009; Tibbetts and Huang, 2010). Indeed, as predicted by the challenge hypothesis, JH titers and CA activity are higher during conflicts associated with unstable social situations under both queenless and queenright conditions (Bloch et al., 2002; Bloch et al., 2000a, 1996; Röseler, 1977; Van Doorn, 1989). The JH – ovary axis is not activated in workers that do not experience intensive agonistic interactions of this kind. For example, JH and oogenesis are low in workers that are placed individually (Cnaani et al., 2002; Duchateau and Velthuis, 1989; Röseler PF., 1990; Sibbald and Plowright, 2015) or in relatively stable queenright conditions. The dominant individuals further inhibit CA activity and JH signaling in subordinate workers (Bloch et al., 2000a, 1996; Bloch and Hefetz, 1999; Shpigler et al., 2010). The consequential elevated JH titers in the dominant bees coordinate many tissues involved in reproduction. These include modulation of the fat body, exocrine glands, and ovaries, and stimulating processes such as vitellogenesis and oogenesis (Amsalem et al., 2013; Shpigler et al., 2014, 2010). Our unpublished RNA sequencing analyses further show that JH regulates the expression of hundreds of genes in the brain, which is consistent with the hypothesis that JH influences behaviors that are associated with reproduction. Thus, JH may stabilize the dominance hierarchy by enhancing the aggressiveness in the most dominant individual, which in turn inhibits its levels in the subordinate ones. It should be noted, however, that currently there is no evidence that JH acts directly on the neuronal and molecular underpinnings of aggression, and its effects may be mediated by additional neuroendocrine signals. For example, dominance in bumble bees is also correlated with neuroendocrine signals such as octopamine (Bloch et al., 2000c), ecdysteroids (Bloch et al., 2000; Geva et al., 2005), and Vg (Amsalem et al., 2014b, 2014a). As for ecdysteroids, although the activated ovaries of the dominant individuals produce high amounts of ecdysteroids (Geva et al., 2005), these probably have little influence (if at all) on aggression and dominance hierarchy formation because ovariectomized workers can attain high dominance rank even if their ovaries are removed at an early age (van Doorn, 1989). Factors other than JH are also important after the initial elevation of JH titers in dominant bees, as evident by the limited effect of JH manipulation in bees that have already established dominance hierarchy in Exp. 4. Additional studies in which the different factors are manipulated are needed for clarifying how and when they interact to regulate aggression and dominance.

### Conclusions

We used multiple experimental designs to manipulate JH in quartets of orphan bumble bee workers to clarify some of the ambiguity concerning the behavioral functions of JH in conflicts over reproductive dominance. We unequivocally show that JH affects aggressiveness and dominance rank in *B. terrestris*, and under some conditions can even be the pivotal factor determining rank during the establishment of a dominance hierarchy. We also show that JH is not the only player in the system. Body size, age, and previous experience are also important and the interplay between these multiple factors is just starting to be unveiled. Our study highlights many open questions pertaining to the mechanisms by which JH affects aggressiveness and social behavior to shape a dominance hierarchy. Although our results show that JH manipulations affect these behaviors, it is yet not clear if JH acts directly on the brain or that its effect is mediated by some yet unknown intermediary neuroendocrine signals.

## Acknowledgements

We thank Michael Schneider and David Zlotkin for helping with behavioral observations. This research was supported by grants from the US-Israel Binational Agricultural Research and Development Fund (BARD, number IS-4418-11, and IS-5077-18, to G.B.) and The Planning and Budgeting Committee (PBC) Fellowship Program for Outstanding Chinese and Indian Post-Doctoral Fellows (to A.P).

## Supplementary figures

**Supplementary Figure S1:**
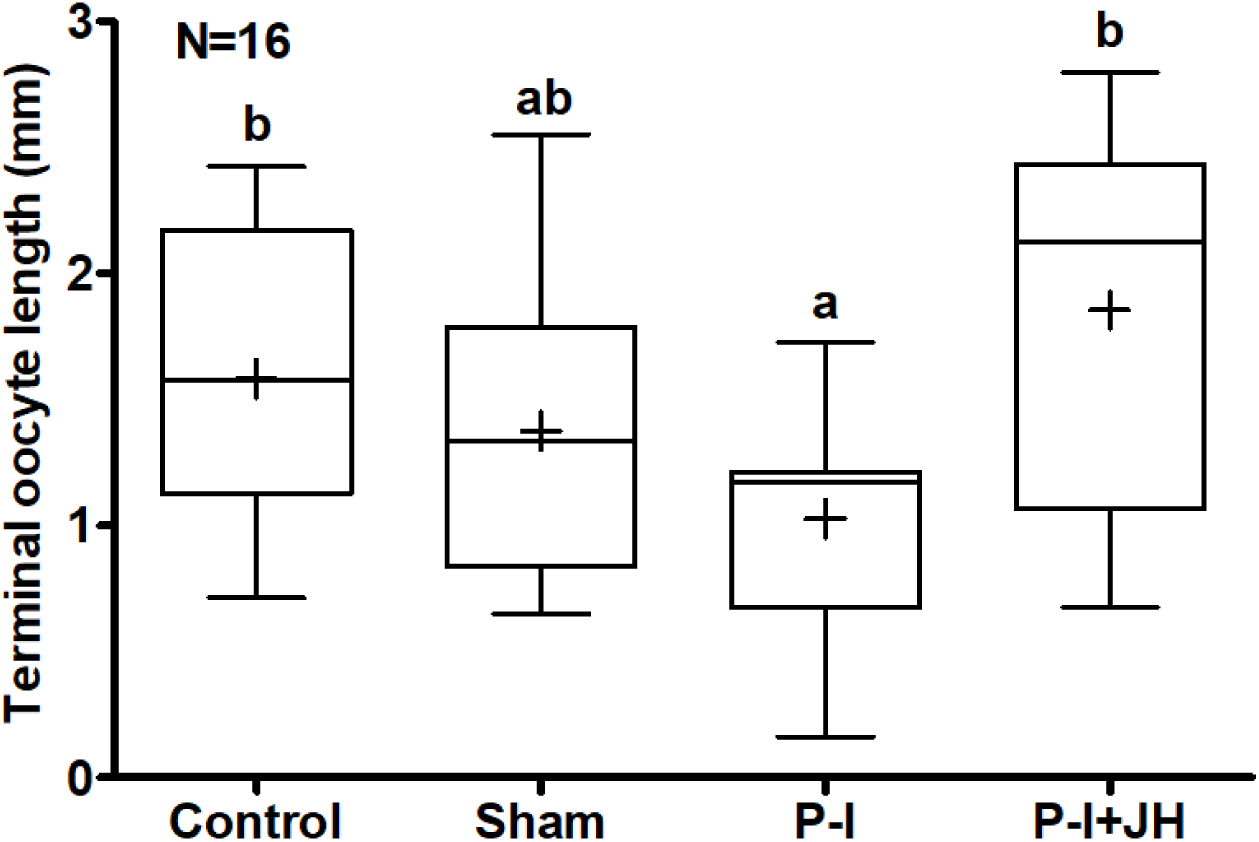
Possible lateral transfer of drugs used to manipulate JH titers. The graph shows ovarian state for bees in a preliminary experiment in which individuals subjected to different treatments were housed together shortly after treatment on Day-1. The ovaries of the P-I treated bees in this experiment are better developed compared with Experiment 1 in which the treated bees where kept individually isolated for the 48hrs after treatment. Treatments marked with different small letters are statistically different in a Kruskal-Wallis test followed by Dunn’s posthoc analysis. Other details as in Figure 1.

**Supplementary Figure S2:**
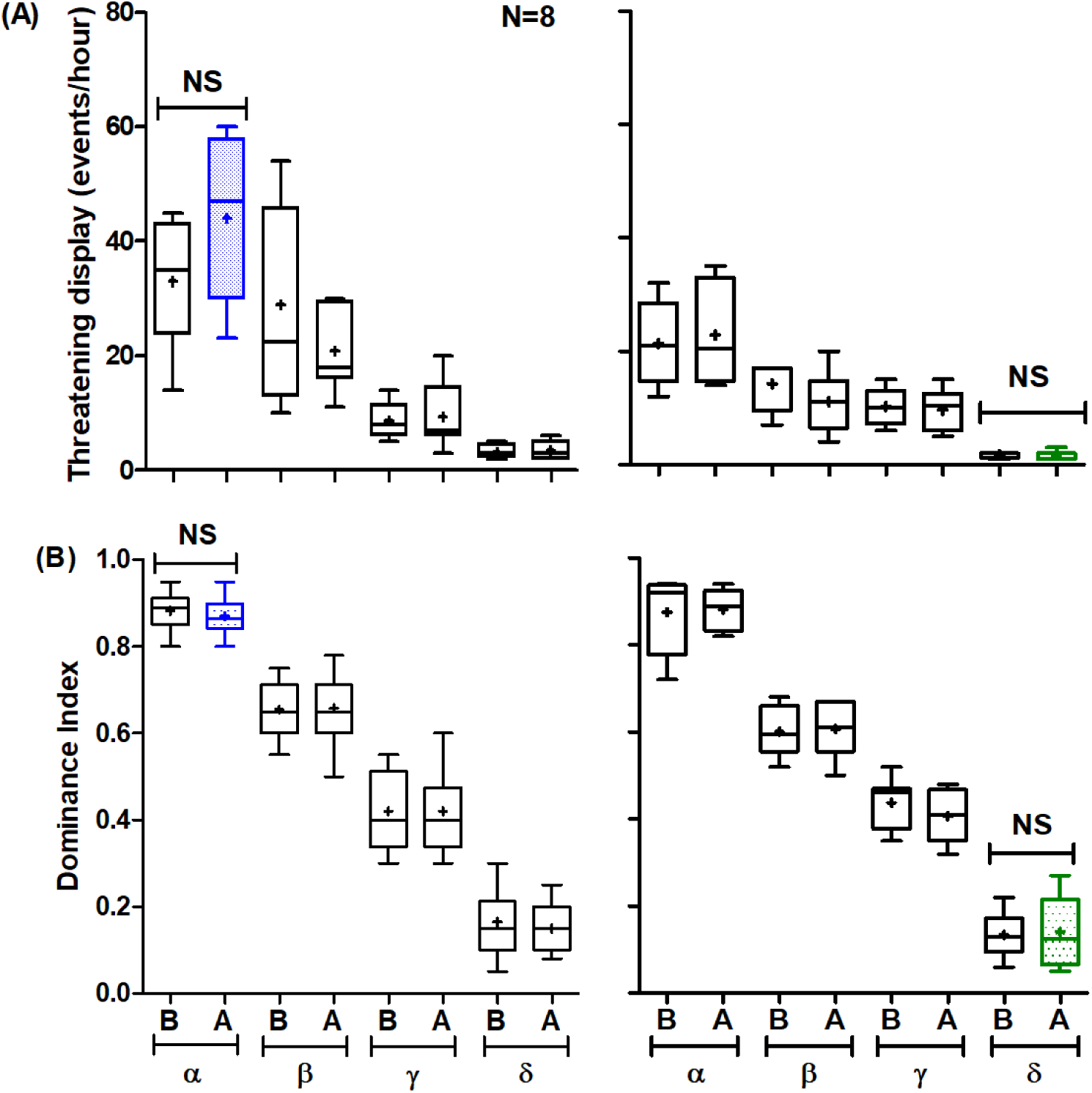
Effect of sham treatments with vehicle only on aggressiveness and dominance rank in groups which had already established stable hierarchies. Left column – Exp. 4A, groups in which we treated the dominant individual with castor oil (5µl/bee, applied to the thorax); right column – Exp. 4B, groups in which we treated the most subordinate individual with DMF (5µl/ bee, applied to the abdomen). (**A**) Threatening behavior and (**B**) Dominance Index B-***Before*** and **A-*After*** the treatment. Other details as in Fig. 1 & 5.

**Supplementary Figure S3:**
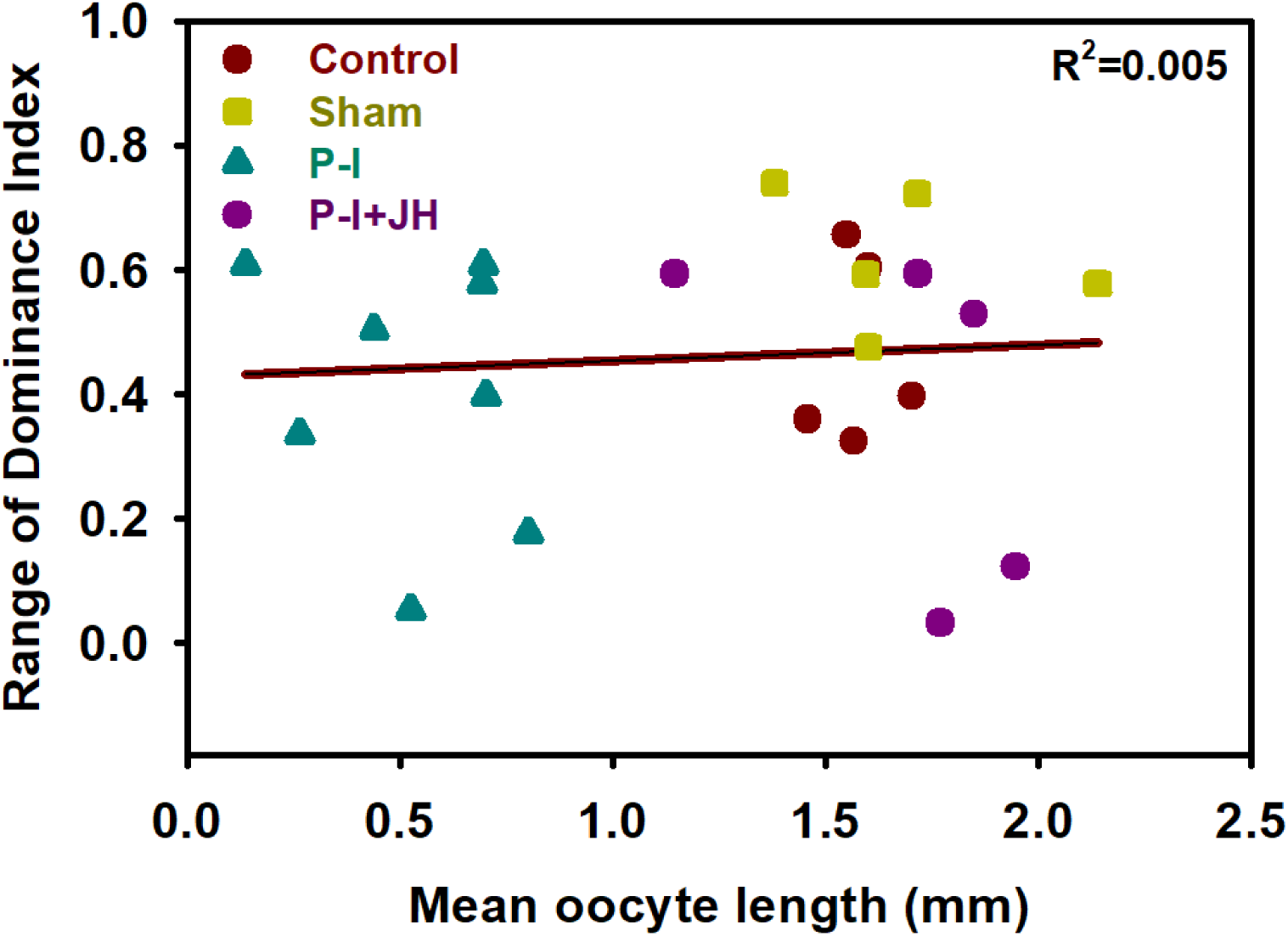
Effect of JH on the linearity of the dominance hierarchy in Experiment 2. The range of Dominance Index is the difference between the Dominance Index of the α and the δ in each group. Treatment groups consisted of different size, similarly-treated queenless worker bees. Other details as in Fig. 4D.

**Supplementary Table S1.** Size-adjusted amounts of Precocene-I and castor oil topically applied to *B. terrestris* workers in Experiments 1-3. The final concentration applied to all the bees was 50µg P-I/ 1µl castor oil.

| Serial Number | Wing size (mm) | Body size denotation | P-I amount (μg/bee) | Castor oil Volume (μl/bee) |
| --- | --- | --- | --- | --- |
| 1 | 2.5-2.6 | Small (S) | 200 | 4.0 |
| 2 | 2.7-2.8 | Small medium (SM) | 220 | 4.4 |
| 3 | 2.9-3.0 | Large medium (LM) | 240 | 4.8 |
| 4 | 3.1-3.2 | Large (L) | 260 | 5.2 |

**Supplementary Table S2.** Correlations between three measures of dominance hierarchy and two measures of oocyte size in the Experiment 2.

| Dependent Variable | Independent Variable | Correlation Coefficient | P value |
| --- | --- | --- | --- |
| Dominance Range | Mean oocyte size | 0.04 | 0.852 |
|  | Maximal oocyte | 0.12 | 0.576 |
| Dominance SD | Mean oocyte size | 0.03 | 0.877 |
|  | Maximal oocyte | 0.14 | 0.515 |
| (Shannon)<sup>-1</sup> | Mean oocyte size | -0.1 | 0.653 |
|  | Maximal oocyte | 0.025 | 0.911 |

**Supplementary Table S3.** Correlations between three estimates of dominance hierarchy and two measures of oocyte size in the Experiment 3.

| Dependent Variable | Independent Variable | Correlation Coefficient | P value |
| --- | --- | --- | --- |
| Dominance Range | Mean oocyte size | 0.63 | < 0.001 |
|  | Maximal oocyte | 0.78 | <10<sup>-5</sup> |
| Dominance SD | Mean oocyte size | 0.66 | < 0.001 |
|  | Maximal oocyte | 0.80 | <10<sup>-6</sup> |
| (Shannon)<sup>-1</sup> | Mean oocyte size | 0.39 | 0.051 |
|  | Maximal oocyte | 0.63 | < 0.001 |

